# Cell fusion upregulates PD-L1 expression and promotes tumor formation

**DOI:** 10.1101/2022.06.14.496068

**Authors:** Youichi Tajima, Futoshi Shibasaki, Hisao Masai

## Abstract

MSCs (mesenchymal stem cells), responsible for tissue repair, rarely undergo cell fusion with somatic cells. Here, we show that approximately 5% of bladder cancer cells (UMUC-3) fuses with bone marrow-derived MSC (BM-MSC) in co-culture and exhibits increased tumorigenicity. Eleven fusion cell clones are established, and 116 genes are identified whose expression is specifically altered in the fusion cells. Many of them are interferon-stimulated genes (ISG), but are activated in a manner independent of interferon. Among them, we show that PD-L1 is induced in fusion cells, and its knockout decreases tumorigenesis in a xenograft model. PD-L1 is induced in a manner independent of STAT1 known to regulate PD-L1 expression, but is regulated by histone modification, and is likely to inhibit phagocytosis by PD1-expressing macrophages, thus protecting cancer cells from immunological attacks. The fusion cells overexpress multiple cytokines including CCL2 that causes tumor progression by converting infiltrating macrophages to tumor-associated-macrophage (TAM). The results present mechanisms of how cell fusion promotes tumorigenesis, revealing a novel link between cell fusion and PD-L1, and underscores the efficacy of cancer immunotherapy.

## Introduction

Cell fusion is a multistep process involving cell-to-cell adhesion, cellular/nuclear membrane remodeling (Hernández & Podbilewicz, 2017). It plays an important role in maintaining biological homeostasis in multicellular organisms, such as osteoclasts or myofibers (Oren-Suissa & Podbilewicz, 2007). During tumor evolution, tumor cells undergo cell fusion to adapt to new micro-environment and to promote their cancerous growth (Weiler & Dittmar, 2019). A number of studies reported that fusion cells are found within a solid tumor and in the circulating blood (Gast Charles *et al*; Yin *et al*, 2020). These fusion cells often exhibit characteristics distinct from the parental cells, including increased tumorigenic potential (Zhou *et al*, 2015).

Mesenchymal stem cells (MSCs) are multipotent stem cells with high self-renewal and differentiation potentials, and are important sources of adult stem cells for cell-based regenerative medicine (Pittenger *et al*, 2019) (Karnoub *et al*, 2007). Bone marrow-derived MSCs (BM-MSCs), in response to tissue damage, migrate to the damaged tissue sites, thereby promoting their repair and cell proliferation by secreting growth factors and molecules assisting tissue regeneration (Alvarez-Dolado *et al*, 2003; Nygren *et al*, 2004; Rizvi *et al*, 2006). Similar to the damaged tissue, BM-MSCs are also recruited to tumor and contribute to immune tolerance and induce angiogenesis necessary for tumor growth (Pawelek & Chakraborty, 2008). The tumor cells then fuse with their surrounding BM-MSCs, frequently resulting in complex aneuploid karyotypes and neoplastic transformation (Johansson *et al*, 2008) (Nygren *et al*, 2008; Rappa *et al*, 2012) (Delespaul *et al*, 2019). However, it is unclear how fusion cells undergo neoplastic transformation. Recently, it was suggested that cell fusion can induce transcriptional reprograming (Feliciano *et al*, 2021).

Cell fusion is also induced by the addition of chemicals or by bacterial or viral infection, and is known to promote the expression of the type I interferon (IFN) (Holm *et al*, 2012; Ku *et al*, 2020). IFN activates IFN-stimulated genes (ISGs), which regulate viral replication and innate immune responses via directly binding to IFN-responsive elements at the promoters (Schoggins, 2019). IFN also upregulates programmed death-ligand 1 (PD-L1, also named CD274), which serves as an ‘immune checkpoint’. PD-L1 binds to receptor programmed death-1 (PD-1, also called PDCD1) to suppress immune responses by antigen presenting cells and T-cells. However, it is not known whether cell fusion induced by non-pathogen stimuli affects immune responses.

In this study, we established hybrid cells (HB cells) between bone marrow-derive mesenchymal stem cell line (BM-MSC) and urothelial carcinoma cell line (UMUC-3) through co-culture. Although the gene expression profile of the HB cells resembles that of BM-MSC, HB cells gained prominent tumorgenicity not seen in the parental cells. In contrast to the parental cells, HB cells constitutively express PD-L1 and the high-level expression of PD-L1 is responsible for their enhanced tumor growth in xenograft mouse model. PD-L1 expression was not regulated by ISGs, but was suppressed by JQ1, an inhibitor of BRD4, epigenetic transcriptional activator. We conclude that cell fusion promotes tumor growth through enhanced expression of PD-L1, that is caused not by IFN induction but by epigenetic modification.

## Results

### Spontaneous cell fusion between MSC and cancer cells

Using the retrovirus-mediated gene transfer, we generated immortalized BM-MSC cells expressing green-fluorescence protein (GFP) and three tumor cell lines, UMUC-3 (human urothelial carcinoma), PANC-1 (pancreatic cancer cells), and MCF-7 (breast cancer cells), each expressing red fluorescent protein (mCherry). The BM-MSC or three tumor cells also expressed neomycin- or puromycin-resistant gene, respectively, that is joined to GFP or mCherry through internal ribosome entry sites (IRES) sequence. To analyze spontaneously-arising fusion cells, we co-cultured BM-MSC-GFP with either UMUC-3-mCherry or PANC-1-mCherry or MCF-7-mCherry at a ratio of 1: 3 for two days, and then analyzed fusion cells by flow cytometry (**Fig. EV1A**). The fusion rates were similar among the three cancer cell lines (1-5 %) (**Fig. EV1B**).

Since the fusion efficiency of UMUC-3 and BM-MSCs was the highest (5%) in the preliminary experiments, they were used to generate stable fusion cells. Co-culture of BM-MSC-GFP and UMUC-3-mCherry in the medium containing both neomycin and puromycin for 9 days generated spontaneously-arising fusion cells in 40% of the population (**Fig. 1A**). We isolated 11 fusion clones (from HB1 to HB11) by limiting dilution. More than 90% of each clone population was GFP- and mCherry-double positive (**Fig. EV1C**). Next, we performed a karyotype analysis of a fusion clone (HB6) and parent cells. BM-MSC contained a near-tetraploid chromosome sets (modal chromosome number 88), and UMUC-3 cells contained a near triploid chromosome sets (modal chromosome number 69) (**Fig. 1B and C**). In contrast, HB6 contained a near-hexaploid chromosome sets (modal chromosome number 142), which is close to the sum of the BM-MSC and UMUC-3 chromosome complements. However, the numbers of chromosomes were distributed over a wide window, ranging from 80 to 159 in most cells with the peak number between 140∼149 (**Fig. 1B and C**). These results indicate that cell fusion generated significant karyotype instability.

**Figure 1.**
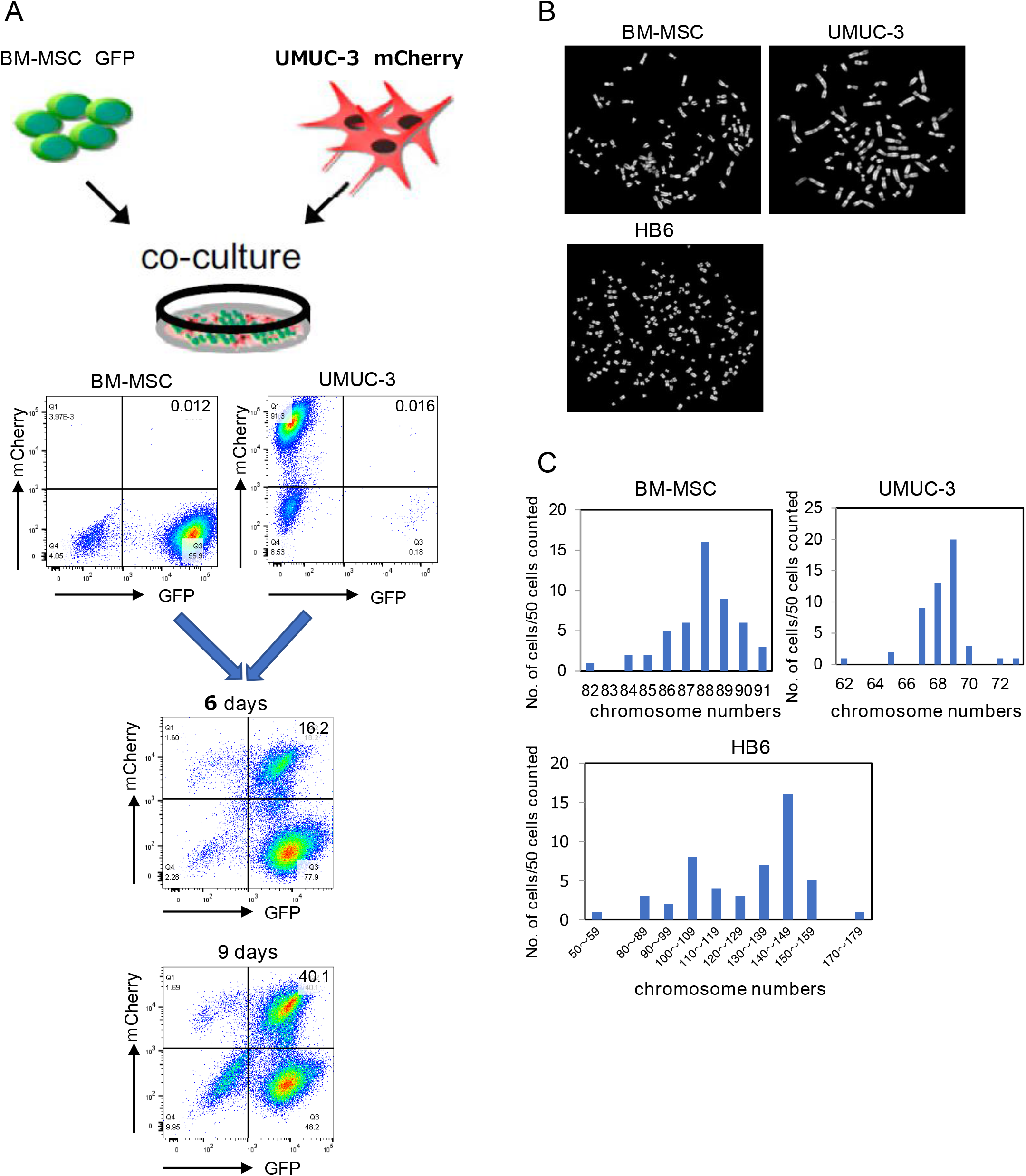
UMUC-3 cells (human urothelial carcinoma) fuse with BM-MSC to form fusion cells with full chromosome complements. **A**. The strategy for selecting fusion cells between UMUC-3 and BM-MSC cells by dual antibiotics selection and isolation of fusion cell clones by limiting dilutions. BM-MSC-GFP spontaneously fused to UMUC-3-mCherry in medium containing puromycin and neomycin. Fusion process was monitored at 3 days, 6 days, and 9 days post-selection by dual color flow cytometry analysis, GFP (FITC-A channel) and mCherry (PE-Texas Red-A channel). Percentages of dual GFP-mCherry positive fusion cells are shown in upper right quadrant of each panel. **B**. Representative metaphase spreads for karyotype analysis of BM-MSC, UMUC-3 and HB6 fusion cell clone. DNA was stained with 4’,6-diamidino-2-phenylindole (DAPI). **C**. Quantification of chromosome numbers in BM-MSC, UMUC-3 and HB6 by karyotyping (*n* = 50 cells for each cell type). Distribution of numbers of cells containing indicated numbers of the chromosome is shown for each cell type.

### Tumorigenic potential of fusion cells (HBs)

We next examined the transcription profiles of eleven HB fusion cells by RNA-seq analyses. Principle component analysis (PCA) of gene expression (**Fig. 2A**) was robustly separated, with 19.5 % and 14.7 % of variance explained in PC1 and PC2, respectively. HB cells, except for HB3, exhibited expression profiles similar to BM-MSC cells rather than to UMUC-3 cells. To investigate the effect of cell fusion on the growth property, we measured the growth rate of fusion cells by a high-content image-based cell counting. The growth rate of HB8 was faster than UMUC-3, whereas that of HB1 and HB7 was slower than UMUC-3 and was similar to that of BM-MSC cells. The growth rate of other HB clones were similar to UMUC-3 cells (**Fig. 2B**).

**Figure 2.**
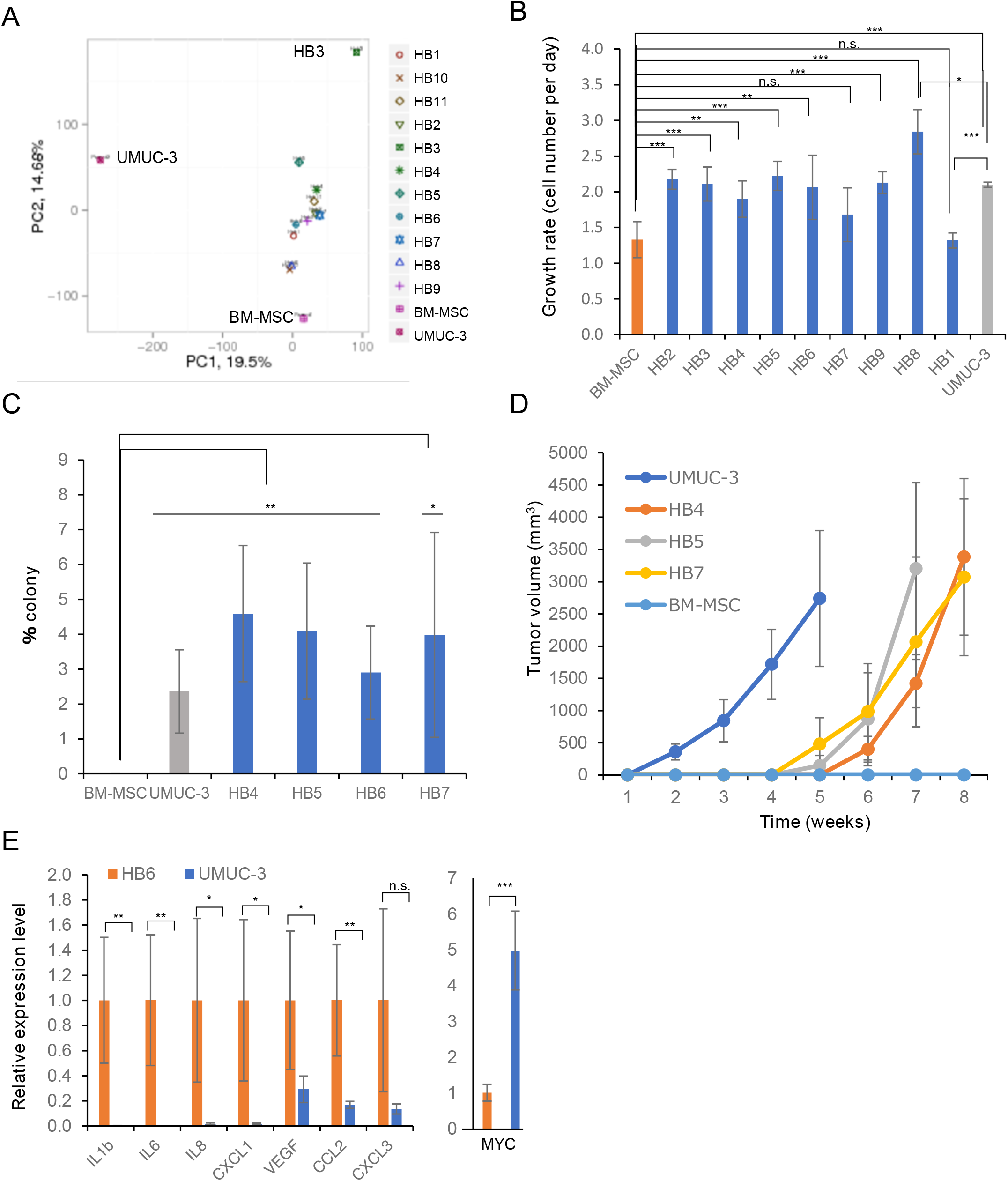
Fusion cells are tumorigenic *in vitro* and *in vivo*. **A**. PCA (Principle component analysis) analysis of BM-MSC, UMUC-3 and HBs according to FPKM (expected number of Fragments Per Kilobase of transcript sequence per Millions base pairs sequenced). **B**. In vitro cell proliferation (mean ±SD) of nine fusion cells, UMUC-3 and BM-MSC as measured by a high content screening (HCS) method. Values are means ± SD. ***, ** and * indicate P<0.001, P<0.01 and P<0.05, respectively, using two-tailed Student’s t-test. **C**. Anchorage-independent clonogenicity in soft agar. Values are means ± SD, ** and * indicates *P*<0.01 and *P*<0.05, respectively, using two-tailed Student’s *t*-test, compared to parental BM-MSC. **D**. Tumor growth of fusion cells, UMUC-3 and BM-MSC after subcutaneous xenograft in immunocompromised nude mice (BALB/c AJcl-Foxn1^nu^). BM-MSC did not lead to tumor growth. UMUC-3, n=15; BM-MSC, n=10; HB4, n=5; HB5, n=5; and HB7, n=5. **E**. Quantitative RT-PCR (RT-qPCR) analysis of cytokines and MYC expression in tumor tissues derived from both HB6 and UMUC-3. Values are means ± SD. ***, ** and * indicate *P*<0.001, *P*<0.01 and *P*<0.05, respectively, using two-tailed Student’s *t*-test.

To evaluate the tumorigenic potential of fusion cells, we first tested the ability of anchorage-independent growth in the soft agar media. Consistent with the growth property *in vitro*, the colony formation rate of BM-MSC was zero, whereas that of fusion cells (HB4, HB5, HB6 and HB7) was 2.5 to 4.5 % compared to 2% with UNUC-3 (**Fig. 2C**).

Three fusion clones (HB4, HB5 and HB7) as well as parent BM-MSC and UMUC-3 were subcutaneously engrafted in immunodeficient nude mice, and the recipient mice were monitored for 8 weeks for formation of tumors. As expected, UMUC-3 cells rapidly induced tumors (take rate of 100%; n=15) (Tanaka & Grossman, 2003), while BM-MSC did not (take rate of 0 %; n=10). Fusion cells, HB4, HB5 and HB7, exhibited tumor growth only at 4 to 5 weeks after transplantation with a take rate of 100% (HB4, n=7; HB5, n=6; HB7, n=6) (**Fig. 2D**). After the tumors started to grow, the rate of tumor growth of HB cells was as fast as that of UMUC-3. HB cells possibly gained the tumorigenic potential by cell fusion with UMUC-3, while maintaining characteristics as mesenchymal stem cells.

We then investigated the mRNA expression of cytokines and growth factors in tumor tissues derived from both HB6 and UMUC-3 by RT-qPCR. Several inflammatory cytokines including IL-1, IL-6, CXCL1, IL-8, VEGFA and CCL2 were more abundantly expressed in tumors from fusion clone HB6 than in those from UMUC-3 (**Fig. 2E**). CCL2 is a monocyte-chemotactic protein and its high level expression correlates with increased numbers of tumor associated macrophage (TAM) in the HB6-derived tumor tissues (Li *et al*, 2017). On the other hand, MYC was expressed more abundantly in UMUC-3 urothelial carcinoma cell-derived tumors (**Fig. 2E**). These results suggest differential tumorigenic properties of HB and UMUC3 cells may be related to distinct transcriptional profiles of these cells.

### Gene expression profile of HB cells in comparison with that of parental cells

MSCs is characterized by expression of CD105, CD73, and CD90 but not that of CD45 or CD34 and by their ability to differentiate into osteoblasts, adipocytes, and chondrocytes *in vitro* (BÜHring *et al*, 2007). The BM-MSC used in this study mostly follow these features (**Fig. EV2A**), bur also strongly expresses α-smooth muscle actin (α-SMA), PDGF receptor-α (PDGFR−α) and vimentin, markers of MSC-derived myofibroblasts and MSC-derived cancer associated fibroblasts (**Fig. EV2A**) (Soliman *et al*, 2021). BM-MSC expressed extracellular matrix components included type 1 collagen (Col1), and fibronectin (FN). Consistent with the RNA-seq results, COL1A1, FN1, and α-SMA proteins were highly expressed in the fusion cells but not in UMUC-3 (**Fig. EV2B and C**). Fusion clones, cultured under adipogenic induction condition, showed marked adipogenic differentiation ability and expressed FABP4, an adipocyte specific marker protein (**Fig. EV2D**). Androgen receptor (AR), expressed and localized to the nucleus in UMUC-3, but not in BM-MSC (**Fig. EV2E**) (Miyamoto *et al*, 2007), is not localized to the nucleus in HB2 and HB8, indicating that some of the gene expression profiles in HB cells have diverged from that of UMUC-3 (**Fig. EV2E**).

To more extensively search for the genes responsible for enhanced tumorigenicity of HB cells, we analyzed transcription profiles of approximately 24,000 genes in HB and the parental cells. We defined upregulated or downregulated genes using cutoff value of 2-fold-change (FC) and false discovery rate (FDR) *q* value < 0.005. Using these criteria, we identified 239 ∼ 499 genes that are differentially expressed between each one of the ten HB cells and BM-MSC, whereas 1335 ∼ 1839 genes were differentially expressed in HB and UMUC-3 (**Fig. 3A**). Yellow, blue, and purple in the Venn-diagrams show differentially expressed genes (DEGs) between HB and BM-MSC, between HB and UMUC-3, and between BM-MSC and UMUC-3, respectively (**Fig. 3A**). We ultimately identified 264 genes differentially expressed between HBs and the parental cells (BM-MSC and UMUC-3 in common) after exclusion of genes differentially expressed between BM-MSC and UMUC-3. Gene ontology analysis revealed that these 264 genes are significantly enriched in 10 biological processes. Interestingly, 33 were involved in response to cytokine (GO:0034097) (**Fig. 3C**) and 16 involved in type I interferon signal pathways (GO:0060337) (**Fig.3 C**). Expression of a panel of genes involved in interferon response, including ISG15, IFI6, IFI27, MX1, MX2, OAS1, OAS2, OAS3, IFIT1, IFIT2, IFIT3 and BST2, increased in HB2, HB4, HB5, HB9 and HB11 cells (**Fig. 3C**). These results suggest that cell fusion between BM-MSC and cancer cells activates cytokine-induced signaling and antiviral responses.

**Figure 3.**
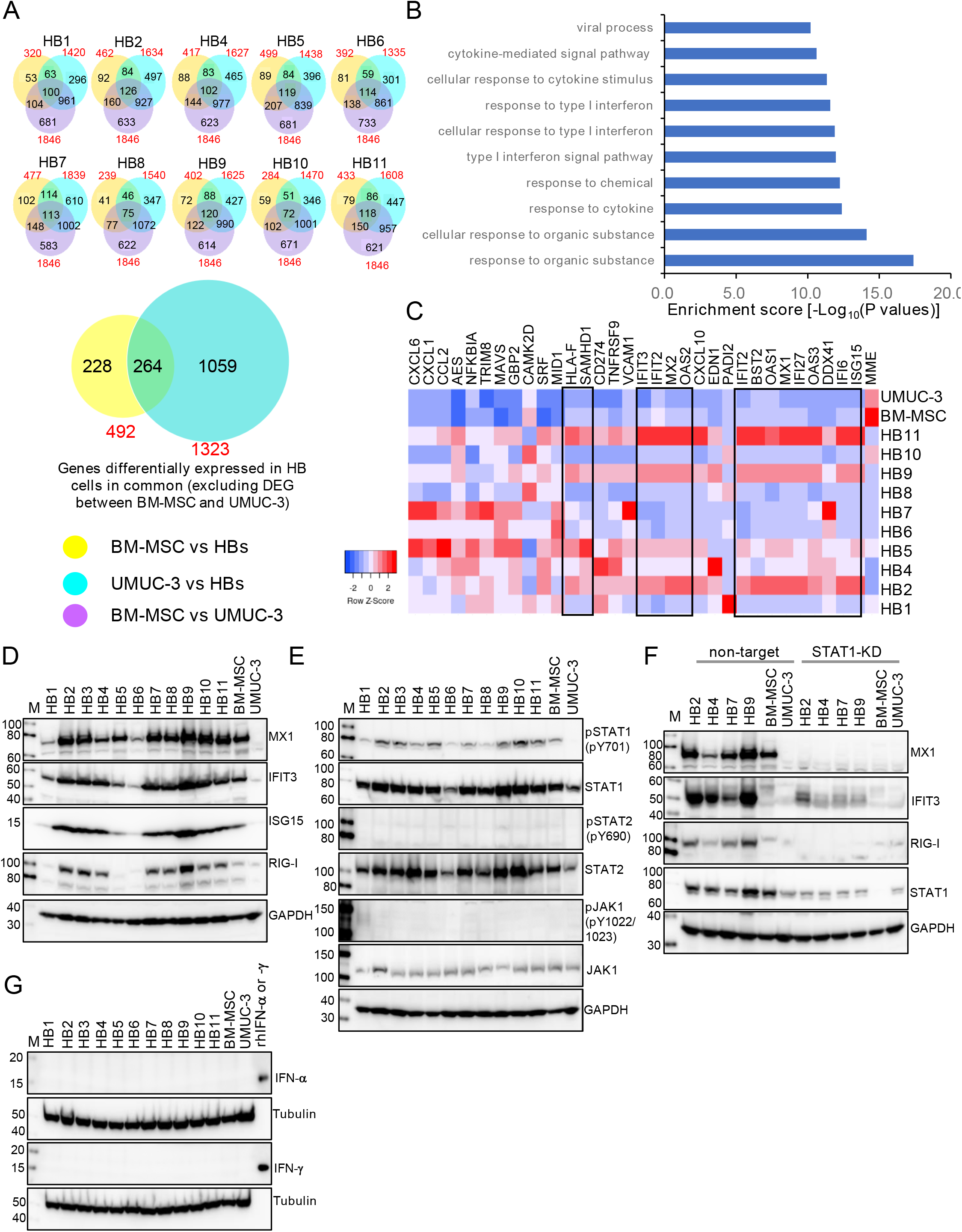
Genes involved in type I interferon signal pathway are activated in fusion cells. **A**. Overlaps of genes differentially expressed between each of the fusion cells (HBs) and BM-MSC (yellow) or UMUC-3 (blue) and those between BM-MSC and UMUC-3 (purple) (upper 10 Venn-diagrams). In total, 264 genes were common among the differentially expressed genes in fusion cells and the two parental cells (after exclusion of differentially expressed genes between BM-MSC and UMUC-3; lower Venn-diagram). **B**. Gene ontology enrichment analysis of the 264 consensus genes. The genes involved in response to cytokine (GO:0034097) and in type I interferon signal pathway (GO:0060337) were among the top enriched gene sets that significantly changed after cell fusion. **C**. Expression levels (Log10 FPKM) of the 33 genes involved in response to cytokine (GO:0034097) are shown as heatmap visualizations by Heatmapper. Sixteen genes involved in type I interferon signal pathway (GO:0060337) are boxed in the heatmap. **D, E** and **G**. Western blot analysis of proteins indicated in parent and eleven fusion cells (from HB1 to HB11). **F**. Western blot analysis of proteins indicated in parent cells and HB2, HB4, HB7, HB9 after STAT1 sgRNA knockdown or control-treatment.

### Cell fusion induces interferon stimulated genes (ISGs) in HB cells

To validate RNA-seq results, we analyzed protein levels by western blotting. Interferon-stimulated genes (ISGs), such as MX1, IFIT3, ISG15and RIG-I, involved in type I interferon response were upregulated in most HB cells, compared with the parental cells (**Fig. 3D**). ISG induction is mediated by activation of the JAK/STAT1 signaling. Interestingly, we found that STAT1 (Y701) is phosphorylated at a significant level in HB cells. However, phosphorylation of JAK1 at Y1022/Y1023 (pJAK1), and that of STAT2 at Y690 (pSTAT2) were not detected (**Fig. 3E**). The knock-down of STAT1 expression by CRISPR/Cas9 led to significant downregulation of ISGs, indicating the role of STAT1 in expression of ISGs in HB cells (**Fig. 3F**). Neither IFN-α nor IFN-γ was expressed in HB cells, excluding the autocrine mechanism of ISGs induction (**Fig. 3G**). These results suggest non-canonical pathway(s) for ISGs induction that involves STAT1 but not the typical type I interferon response pathway.

### Cell fusion results in increased PD-L1 expression

RNA-seq analysis revealed upregulation of programmed death ligand 1 (PD-L1, also named CD274) in HB cells compared with the parental cells. We examined surface PD-L1 expression by immunostaining. PD-L1 staining signals in the fusion clone (HB6) were much higher than those in BM-MSC and UMUC-3 (**Fig. 4A**). The specificity of the PD-L1 antibody used was verified by the absence of the signals in HB6 derived PD-L1 KO cells constructed by gene editing.

**Figure 4.**
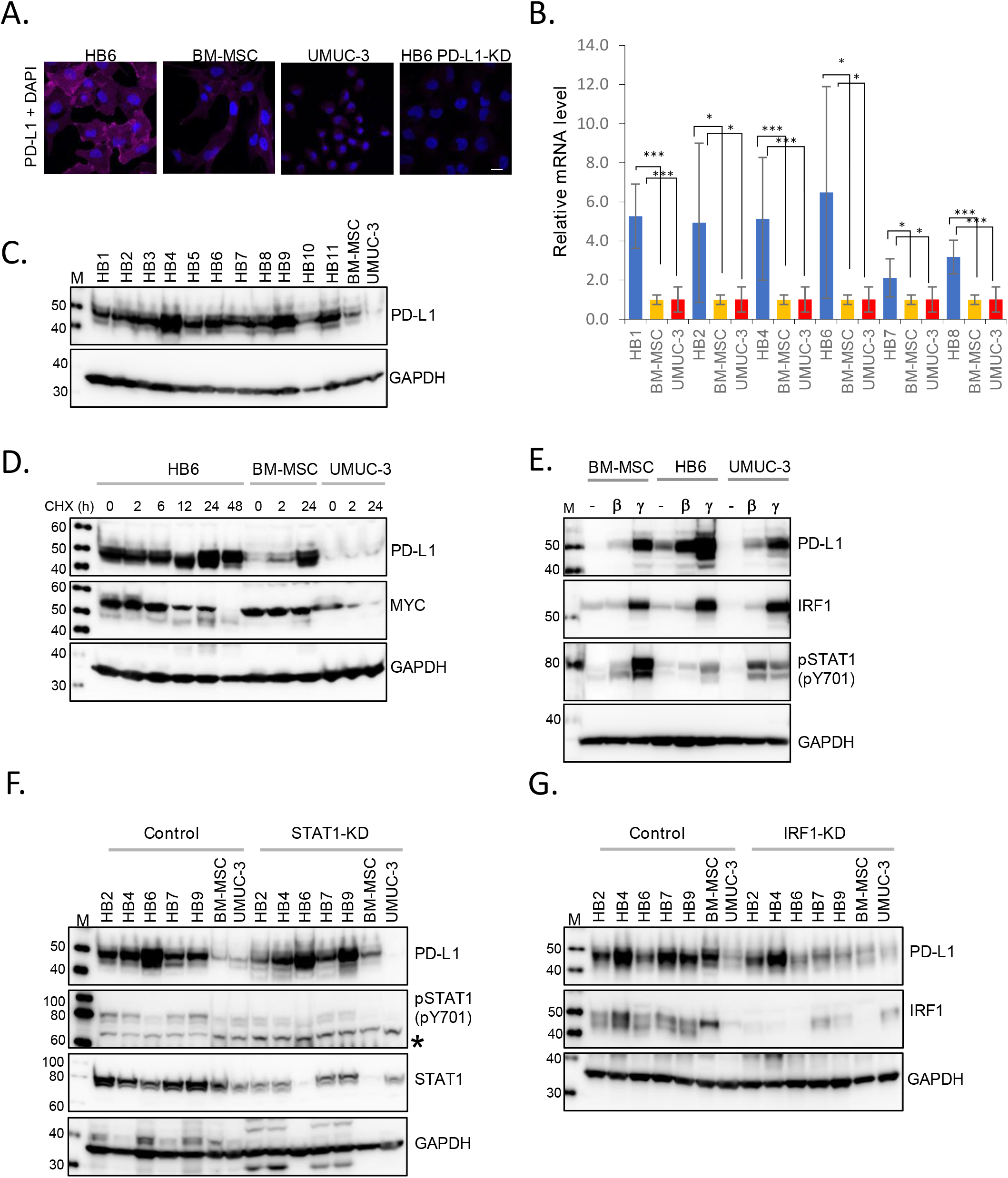
Cell fusion-induced upregulation of PD-L1 expression. **A**. Immunostaining of PD-L1 in parental cells (BM-MSC and UMUC-3), fusion cells (HB6) and HB6 after PD-L1 sgRNA knockdown (KD) (as a negative control). Nuclei were counterstained with DAPI. Magnification, x400. Scale bar = 50 µm. **B**. The mRNA levels of *PD-L1* were determined in parental cells (BM-MSC and UMUC-3) and fusion cells (HB1, HB2, HB4, HB6, HB7 and HB8) by RT-qPCR. The values were normalized to that of ubiquitin C (*UBC*). Each value represents the mean (± SD) from at least three independent experiments. *** indicates *P* < 0.001, * indicates *P* < 0.05 using two-tailed Student’s *t*-test. **C**. Western blot analysis of PD-L1 in parental cells (BM-MSC and UMUC-3) and eleven fusion cells (from HB1 to HB11). **D**. Western blot analysis of proteins indicated in parental (BM-MSC and UMUC-3) and fusion cells (HB6) treated with 100 µg/ml cycloheximide (CHX) for indicated times. It is not clear why the PD-L1 protein level increased at 24 h in BM-MSC. **E**. Western blot analysis of proteins indicated in parental (BM-MSC and UMUC-3) and fusion cells (HB6) treated with IFN-β (β; 100 ng/ml) or IFN-γ (γ; 100 ng/ml) for 24 h or non-treated. **F** and **G**. Western blot analysis of proteins indicated in parent (BM-MSC and UMUC-3) and HB2, HB4, HB6, HB7, HB9 after non-target sgRNA and STAT1 (**F**) or IRF1 (**G**) sgRNA. In F, the asterisk indicates a nonspecifically reacting band.

We detected significant upregulation of PD-L1 at both RNA and protein levels in HB cells through RT-qPCR and western blotting analyses, respectively (**Fig. 4B and C**). Since PD-L1 protein is known to be subject to the ubiquitin-mediated proteasomal degradation (Burr *et al*, 2017a; Mezzadra *et al*, 2017; Zhang *et al*, 2018a), we treated HB6 and parental cells with cycloheximide (CHX) to block *de novo* protein synthesis and examined the protein stability. Unlike the results from previous studies (Li *et al*, 2016a), the PD-L1 protein did not change for 48 h after addition of CHS in HB6, whereas MYC protein decreased (**Fig. 4D**). Both PD-L1 and MYC proteins were unstable in UMUC-3 cells under the same condition (**Fig. 4D**). These results indicate that PD-L1 expression is significantly upregulated by cell fusion through multiple mechanisms.

### Constitutive expression of PD-L1 in HB cells is independent of STAT1 pathways

PD-L1 expression can be induced by IFN-α, -β, and -γ in a JAK1/2-STAT1/2-IRF1-dependent manner (Garcia-Diaz *et al*, 2019). Consistent with this, we found that IFN-γ increased PD-L1 expression, concomitant with upregulation of IRF1 and STAT1 phosphorylation, in HB6, UMUC3, and BM-MSC cells (**Fig. 4E**). To examine the molecular mechanism of constitutive PD-L1 expression in HB cells, we examined the effects of knock-down of the genes involved in IFN signaling pathway on the basal level of PD-L1 expression. We have knocked-down STAT1 and IRF1 in HB cells by CRISPR/Cas9-mediated genome editing technology. We confirmed the depletion or significant downregulation of *STAT1* and *IRF1* proteins by Western blotting (**Fig. 4F and 4G**). Although the knock-down of STAT1 did not significantly affect the expression level of PD-L1 (**Fig. 4F**), the depletion of IRF1 reduced the expression level of PD-L1 (**Fig. 4G**). These results indicate that increased expression of PD-L1 in HB cells is independent of STAT1 signaling pathway but partially depends on IRF1.

### High-level expression of PD-L1 in fusion cells is suppressed by BRD4 inhibitor, JQ1

RNA-seq analyses suggests that expression profiles of key epigenetic modifiers, such as SMARCC1 (BAF155) and SMARCB1 (SNF5), significantly changed in HB cells compared to the parental cells, implying that cell fusion causes dramatic epigenetic changes (**Fig. 5A**). The bromodomain and extraterminal (BET) protein BRD4 functions as an epigenetic reader via direct binding to acetylated lysine on histone, thereby activating transcription. To examine the involvement of BRD4 in PD-L1 expression, we treated HB cell with JQ1, a potent BRD4 inhibitor, and examined the expression of PD-L1. JQ1 decreased the PD-L1 mRNA level in HB cells by more than three to ten fold (**Fig. 5B**), whereas it did not significantly affect the PD-L1 mRNA in BM-MSC and UMUC-3 cells. The PD-L1 protein level in HB cells were also downregulated by JQ-1 (**Fig. 5C**). These results indicate that constitutive and increased PD-L1 expression in HB cells may involve BRD4-mediated epigenetic modification.

**Figure 5.**
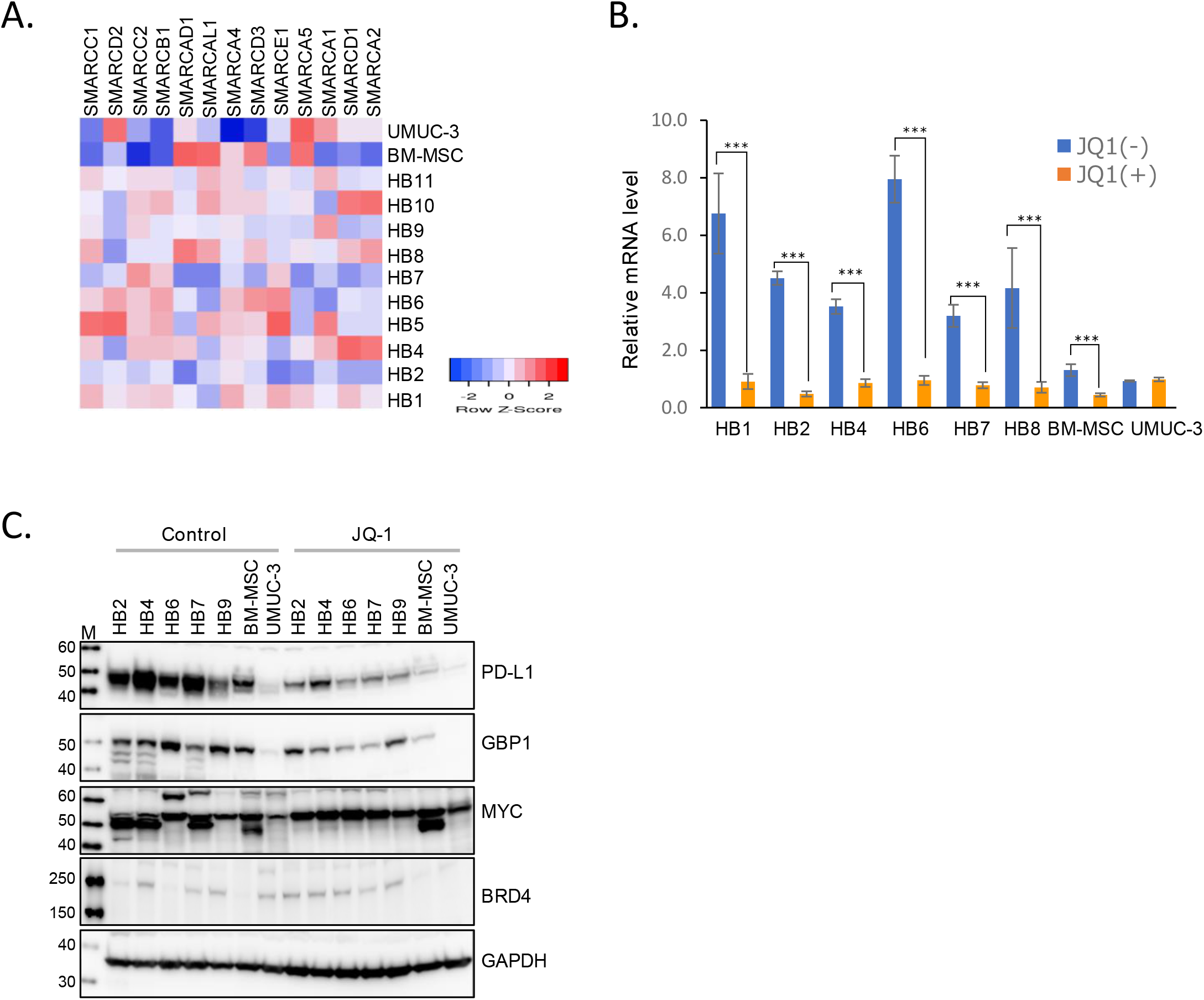
PD-L1 expression in fusion cells is reduced by inhibition of BRD4. **A**. Expression levels (Log10 FPKM) of the 13 genes involved in chromatin remodeling BAF (SWI/SNF) complex in parental cells (BM-MSC and UMUC-3) and HBs are shown as heatmap. **B**. The mRNA levels of *PD-L1* were determined by RT-qPCR in parental (BM-MSC and UMUC-3) and fusion cells (HB1, HB2, HB4, HB6, HB7 and HB8) treated with 1 µM JQ-1 for 24 h or not treated. The values were normalized to that of ubiquitin C (*UBC*). Each value represents the mean (± SD) from at least three independent experiments. *** indicates *P* < 0.001, using two-tailed Student’s *t*-test. **C**. Western blot analysis of proteins indicated in parent (BM-MSC and UMUC-3) and fusion (HB2, HB4, HB6, HB7, HB9) cells treated with 1 µM JQ-1 for 24 h or non-treated (control).

### Tumorigenicity of fusion cells are reduced by PD-L1 knockdown in xenograft model

The increased PD-L1 expression in fusion cells may promote tumorigenesis by suppressing cellular immune surveillance. Therefore, we examined the effects of PD-L1 knockdown on the ability of fusion cells to support tumor growth *in vivo*. To this end, we deleted PD-L1 genes using a lentiCRISPRv2-sgRNA targeting PD-L1 in HB6, UMUC3, and BM-MSC cells. We checked PD-L1 deletion by western blotting analysis and confirmed down regulation of PD-L1 in the resulting candidate PD-L1^-/-^ cells (**Fig. 6A**). HB6-derived PD-L1^+/+^ and PD-L1^-/-^ cells were implanted subcutaneously into the flank of immunodeficient nude mice, and tumor development was monitored. Tumor volume of HB6 PD-L1^+/+^ reached 1700 mm^3^ (n=8) at 9 weeks, whereas that of HB6 PD-L1^-/-^ only 490 mm^3^ (n=6), indicating a critical role of PD-L1 expression in tumor growth (**Fig. 6B**).

**Figure 6.**
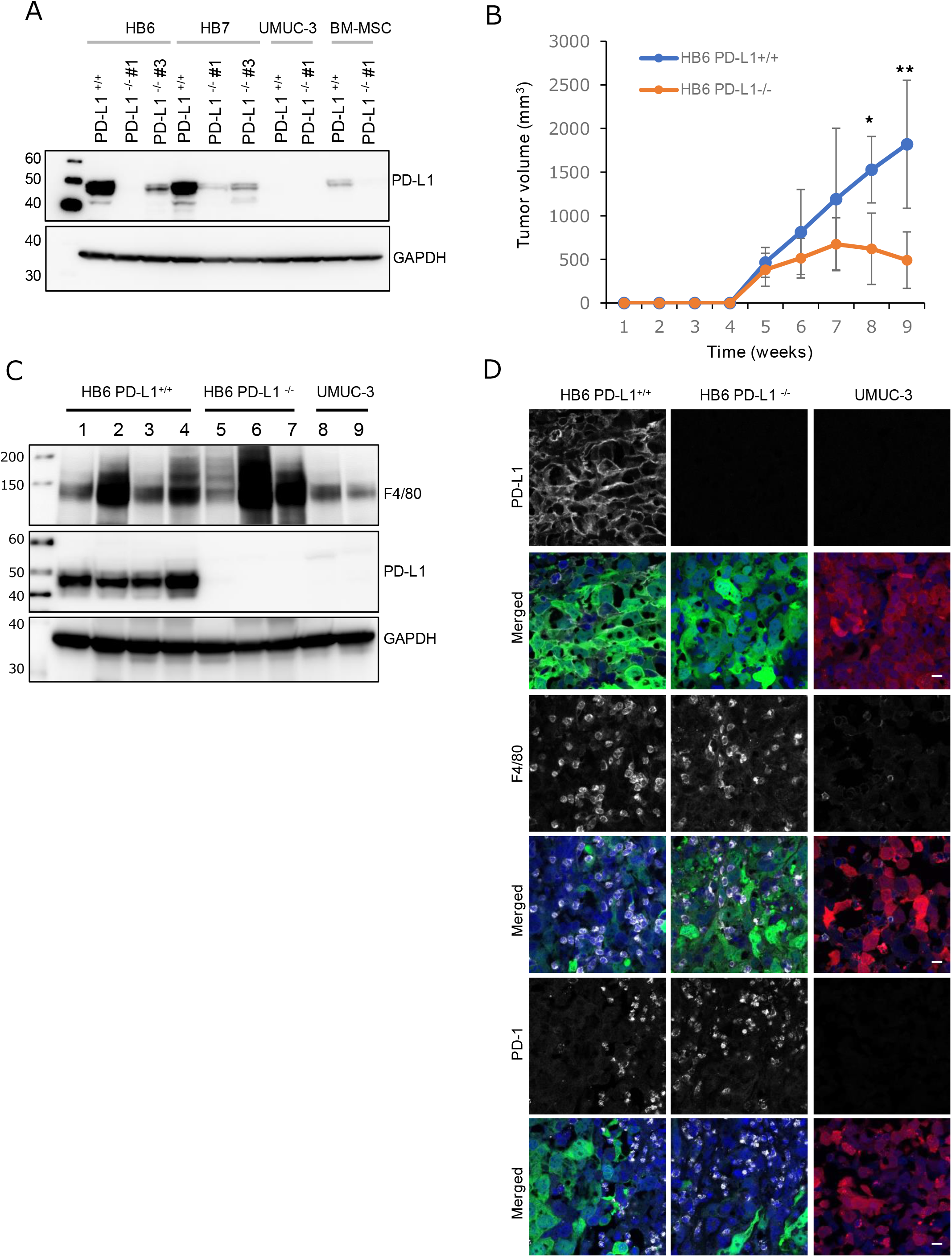
PD-L1 knockdown in the fusion cells results in decreased tumorigenicity in the xenograft model. **A**. Knockdown of PD-L1 in parent (BM-MCS and UMUC-3) and fusion cells (HB6 and HB7). Parent cells and fusion cells were transduced with non-target sgRNA (+/+) and PD-L1 sgRNA (-/-#1 or -/-#3) lentivirus particles. Western blot analyses of PD-L1 and GAPDH in each cell line indicated. **B**. Tumor growth of HB6 with knockdown of PD-L1 (HB6 PD-L1^-/-^; n=6) and that of control HB6 (HB6 PD-L1^+/+^; n=8) in immunocompromised nude mice. **C**. Western blot analysis of proteins indicated in tumors from HB6 PD-L1^+/+^ (4 independent samples, lanes 1-4), HB6 PD-L1^-/-^ (3 independent samples, lanes 5-7) and UMUC-3 (2 independent samples, lanes 8-9). **D**. Immunohistochemistry of PD-L1, F4/80, and PD-1 in tumor tissues derived from HB6 PD-L1^+/+^, HB6 PD-L1^-/-^,and UMUC-3. GFP (green) and mCherry (red) were detected in HB6 and UMUC-3, respectively. Nuclei were counterstained with DAPI (blue). Magnification x400. Scale bar indicate 50 µm.

### Roles of PD-L1 in promotion of tumorigenesis by HB

To examine the infiltrating immune cells, we excised the tumors and performed western blot and immunohistochemical (IHC) staining using anti-mouse macrophage marker F4/80 and anti-mouse PD-1. Our xenograft model was based on athymic nude mice (nu/nu) which is incapable of producing mature T-cells (Pelleitier & Montplaisir, 1975). Tumor tissues from the both PD-L1^+/+^ and PD-L1^-/-^ HB6 cells were infiltrated by F4/80-positive and PD-1-positive mouse macrophages, whereas those from UMUC-3 cells showed significantly reduced infiltration of the same macrophages (**Fig. 6C and D**). We could not find any differences in infiltrating mouse macrophages between tumors from PD-L1^+/+^ and PD-L1^-/-^ HB6cells. The results may suggest that the increased tumorigenic activity of PD-L1^+/+^ fusion cells may be due to suppression of macrophage-mediated phagocytosis through PD-1/PD-L1 immune checkpoint pathway (Gordon *et al*, 2017).

## Discussion

Tumor cells are known to fuse with other cells in the process of their development, and the cell fusion may lead to increased malignancy by promoting infiltration and metastasis. Although cell fusion could affect tumorigenic potential of cancer cells in many ways, the detailed mechanisms are still not clear.

Bone marrow-derived mesenchymal stem cells (BM-MSC) are abundant in tumor environment and play critical roles during tumor progression. Fusogenic BM-MSC are reported to be involved in carcinogenesis, tumor heterogeneity and acquisition of stem cell-like and metastatic properties (Pawelek & Chakraborty, 2008). In this study, we demonstrated that spontaneous fusion between human urothelial carcinoma cell line UMUC-3 and immortalized BM-MSC can efficiently generate hybrid cells. We have discovered that the fusion cells ubiquitously overexpress PD-L1, and that elevated PD-L1 is responsible for tumor promoting activity of fusion cells.

### Increased PD-L1-PD1 interaction may enhance immune evasion of tumor formation by the fusion cells

Tumor tissues from fusion cells were extensively infiltrated by F4/80 positive mouse macrophages, whereas those from UMUC-3 cells exhibited much reduced infiltrating F4/80 positive macrophages (**Fig. 6D**). Multiple cytokines, including IL-1α, IL-6, IL-8, CXCL1, VEGF, CCL2, and CXCL3, are expressed at a higher level in tumors derived from the fusion cells than in UMUC-3 tumors. CCL2 exerts its tumor-promoting effects mainly through recruiting monocytes, contributing to the elevated TAMs that are involved in almost every stage of cancer development (Qian *et al*, 2011). IL-6 has also been shown to promote cell proliferation and enhance tumor metastasis and is required for tumor-initiating cell maintenance via the autocrine IL-6/LIN28 pathway (He *et al*, 2013). Thus, elevated expression of cytokines appears to be a mechanism for the tumor promoting property of the fusion cells.

The enhanced ability of immune evasion may also explain how the fusion cells promote tumorigenesis. We showed that tumor formation of the fusion cells in xenograft is reduced by knockout of PD-L1 (**Fig. 6D**). PD-1 positive macrophages were detected in both PD-L1^+/+^ and PD-L1^-/-^ HB6 tumor tissues. PD-1, first discovered as a coinhibitory receptor on activated T cells, was later shown to be expressed in B cells, NK cells, dendritic cells, monocytes, and macrophages (Gordon *et al*., 2017). Gordon and colleagues (Gordon *et al*., 2017) showed that PD-1/PD-L1 interaction plays suppressive role in macrophages, inhibiting phagocytosis. Upregulated PD-L1 on tumor cells may suppress macrophage-mediated phagocytosis through PD-1/PD-L1 interaction. In fusion cells, expression of CD47, which inhibits the phagocytotic activity of TAM, is induced, which may also contribute to enhanced tumorigenicity.

### How does cell fusion between MSC and cancer cell lead to increased PD-L1 expression?

We observed a 5-fold upregulation of PD-L1 mRNA levels in the fusion cells compared to the parent BM-MSC. PD-L1 transcription in cancer cells is regulated by a variety of transcription factors including HIF-α, NF-κB, IRF1, STAT1, and STAT3 (Sun *et al*, 2018a). However, STAT1 knockdown did not significantly affect the expression of PD-L1, while that of IRF1 partially reduced PD-L1 protein level in the fusion cells (**Fig. 5A**), suggesting that the canonical STAT1/3-IRF1 pathway is not involved in the induction of PD-L1 expression. On the other hand, inhibition of PD-L1 expression by JQ-1, a BRD4 inhibitor, suggests epigenetic regulation of PD-L1 expression. Chromatin may become open at the promoter locus in a manner dependent on histone acetyltransferase BRD4. Further studies are needed to evaluate this possibility.

Since the PD-L1 protein level increases by much greater extent, PD-L1 expression may be regulated at both transcription and post-transcription levels. Indeed, the PD-L1 protein appears to be very stable in HB cells (**Fig. 4D**). Several modifications of PD-L1, including phosphorylation (Li *et al*, 2016b), glycosylation (Li *et al*, 2018; Li *et al*., 2016b), and ubiquitination (Lim *et al*, 2016; Zhang *et al*, 2018b), have been reported to regulate PD-L1 stability. Li-Chuan and colleagues (Chan *et al*, 2019) show that IL-6-activated JAK1 phosphorylates Tyr112 of PD-L1, which recruits the ER-associated N-glycosyltransferase STT3A to promote PD-L1 glycosylation and maintain PD-L1 stability. Thus, increased cytokine expression in the fusion-derived tumors can enhance the posttranslational stability of PD-L1.

Furthermore, PD-L1 is polyubiquitinated in ER. Mezzadra and colleagues (Mezzadra *et al*., 2017) and Burr and colleagues (Burr *et al*, 2017b) identified CMTM6 as a PD-L1 regulator. RNA-seq analysis showed that expression of some proteasome subunit genes including PSMA5, PSMA7, PSMB1, PSMB6 and PSMB7 is reduced in the fusion cells compared to the parent UMUC-3. It is likely that ER-bound PD-L1 is degraded by the 26S proteasome, and thus, reduced proteasome in the fusion cells may be responsible for the increased level of PD-L1. ATR inhibition was reported to destabilize PD-L1 mainly in a proteasome-dependent manner (Sun *et al*, 2018b). Accumulating evidence points to extensive regulation of PD-L1 by the ubiquitin/proteasome pathway (Mezzadra *et al*., 2017), suggesting that targeting of PD-L1 polyubiquitination is an attractive approach to enhanced immune checkpoint therapy.

On the basis of the results presented here, we propose that cell fusion between cancer and mesenchymal stem cells would lead to increased expression of PD-L1 through epigenetic alteration, which would increase its interaction with PD-1 on the infiltrating macrophages, resulting in inhibition of phagocytosis and eventually enhanced tumorigenesis.

## Materials and Methods

### Cells and Reagents

UMUC-3, PANC-1, MCF-7, (all originally from ATCC) were maintained in Dulbecco’s modified Eagle’s medium (DMEM) (Nacalai tesque, Japan) supplemented with 10% (v/v) fetal calf serum (FCS), 100 units/ml penicillin, 100 mg/ml streptomycin (Gibco Life technologies). Human mesenchymal stem cells derived from bone marrow (BM-MSC) was obtained from Cellular Engineering Technologies (CET Inc., Coralville, IA, USA). Immortalized BM-MSC was spontaneously isolated and was cultured in DMEM supplemented with 10% fetal bovine serum, 100U/ml penicillin and 100 mg/ml streptomycin (Invitrogen Life Technologies) in a humidified atmosphere with 5% CO_2_ at 37°C. Antibodies used for immunostaining, flow cytometry analysis, and western blot analyses were as follows. Anti-alpha smooth muscle actin (ab5694, Abcam), anti-collagen I alpha 1 (NB600-408, Novus Biologicals), anti-Fibronectin (15613-1-AP, proteintech), PE/Cy7 anti-human CD140α (PDGFRα) (Clone 16A1, 323507, Biolegend), APC-human CD73 (Ecto-5’-nucleotidase) (clone AD2, 344005, Biolegend), APC/Cy7 anti-human CD90 (Thy1)(clone 5E10, 328131, Biolegend), APC-anti-human CD105 (clone 43A3, 323208, Biolegend), anti-androgen receptor (#5153, Cell Signaling Technology), APC-Human CD34 (clone 4H11, 17-0349, eBioscience), APC-anti-human/Mouse CD44 (Clone IM7, 17-0441, eBioscience), APC-anti-Human CD45 (clone 2D1, 368511, Biolegend), anti-Vimentin (#9856, Cell Signaling Technology), anti-STAT2 (16674-1-AP, proteintech), anti-phospho-Stat2 (Tyr690) (#88410, Cell Signaling Technology), anti-MX1 (13750-1-AP, proteintech), anti-IFIT3 (15201-1-AP, proteintech), anti-Interferon-α (#3110, Cell Signaling Technology), anti-Interferon-γ (15365-1-AP, proteintech), anti-STAT1 (#14994, Cell Signaling Technology), anti-phospho-Stat1 (Tyr701) (#9167, Cell Signaling Technology), anti-USP18 (#4813, Cell Signaling Technology), anti-ISG15 (#2758, Cell Signaling Technology), anti-FABP4 (#2120, Cell Signaling Technology), anti-c-Myc (#5605, Cell Signaling Technology), anti-PD-L1 (GTX104763, GeneTex), anti-PD-1 (mouse specific)(#84651, Cell Signaling Technology), anti-PD-L1 (Extracellular domain specific) (#86744, Cell Signaling Technology), anti-F4/80 (#30325, Cell Signaling Technology), anti-Rig-I (#3743, Cell Signaling Technology),anti-IRF-1 (#8478, Cell Signaling Technology), anti-Jak1 (#3344, Cell Signaling Technology), anti-phospho-Jak1-Y1022/1023 (AP0530, ABclonal), anti-GBP1 (clone 1B1, sc-53857, Santa Cruz Biotechnology), anti-BRD4 (clone C-2, sc-518158, Santa Cruz Biotechnology), anti-GAPDH (Clone 6C5, sc-32233, Santa Cruz Biotechnology), anti-L Tubulin (clone B-5-1-2, sc-23948, Santa Cruz Biotechnology), Alexa Flour 647-linked antibody (#613407; Biolegend), Cy5-linked goat anti-rabbit IgG (111-175-144, Jackson Immuno Research Laboratories), Cy5-linked goat anti-mouse IgG (115-175-146, Jackson Immuno Research Laboratories, Peroxidase-linked goat anti-rabbit IgG, (111-035-144, Jackson Immuno Research Laboratories) and Peroxidase-linked goat anti-mouse IgG (115-035-146, Jackson Immuno Research Laboratories).

### Cell fusion and selection of fusion cells

BM-MSC was generated by retroviral infection using pQCXIP (Clontech Laboratories, Inc., USA) with GFP gene sequence, whereas cancer cell lines were transduced by the retrovirus vector pQCXIN carrying the mCherry gene. Hybrid were made as follows: 2 × 10^5^ cells of BM-MSC-GFP cells and 6 × 10^5^ cells of cancer cell lines-mCherry were seed together in 10 cm dishes. Spontaneous hybrid cells were observed after 2 days by flow cytometry using LSRFortessa flow cytometer (BD Bioscience). To obtain the stable spontaneously fusion cells, 2 × 10^5^ BM-MSC-GFP/neo cells were mixed with 6 × 10^5^ UMUC-3-mCherry/puro cells for 2 days without antibiotics. After 2 days, the coculture was selected with 1 µg/ml of puromycin (Fujifilm-Wako pure chemical Corp.) and 250 µg/ml of neomycin (Nacalai tesque, Japan) to obtain dual resistant cells, or subjected to flow cytometry to purify GFP/mCherry double-positive cells. To isolate fusion cell clones, selected GFP/mCherry double-positive cells were cloned by limiting dilutions. To validate isolated fusion cell clones, they were analyzed by flow cytometry to examine expression of GFP and mCherry.

### Karyotype analysis

Chromosome spreads were prepared, using standard protocols, from parental and fusion cells (HB6) treated with colcemid 100 ng/ml (Fujifilm-Wako pure chemical Corp.) for more than 2 h to induce mitotic arrest. Subsequently, the cells were detached from plates in 0.25% trypsin and treated with hypotonic solution (0.075M KCl) for 15 min. Cells were then fixed by slow dropwise addition of fresh Carnoy’s fixative (methanol: acetic acid 3:1). After washes with the fixative solution 3 times, a drop of the cell suspension was placed on a humidified glass side. Sides were mounted using ProLong™ Gold antifade mountant with DAPI (Thermo Fisher Scientific). The prepared chromosome spreads were analyzed using 60x water objective lens on BZ-9000 fluorescent microscope with appropriate filters (Keyence, Japan). Chromosome numbers were determined for 50 metaphase spreads for each cell line.

### RNA sequencing analysis

The library was prepared for RNA-sequencing using the standard Illumina protocols, and Novogene (Beijing, China) conducted RNA-sequencing using an Illumina HiSeqTM2500 sequencer. Sequencing reads were aligned to the human genome (Hg38) using TopHat2 (Kim *et al*, 2013) (http://ccb.jhu.edu/software/tophat.). The abundance of transcript reflects gene expression level. In RNA-seq experiment, we estimate gene expression level by the abundance of transcripts that mapped to the genome or exon. Reads counts are proportional to gene expression level, gene length and sequencing depth. FPKM (short for the expected number of Fragments Per Kilobase of transcript sequence per Millions base pairs sequenced) is the most common method of estimating gene expression levels, which takes the effects of both sequencing depth and gene length on counting of fragments into consideration. Principal Component Analysis (PCA) of samples were conducted according to FPKM of all samples. Then, make linear combination of these principal components as new composite indicator to do clustering analysis for samples. Differentially expressed gene (DEG) analysis of the 2 parent cells and the 11 fusion cells were performed using the DESeq2 R package (1.10.1) (Love *et al*, 2014) (http://www.bioconductor.org/packages/release/bioc/html/DESeq2.html.). DESeq2 provides statistical routines for determining differential expression in digital gene expression data using a model based on the negative binomial distribution. The *P* -values were adjusted using the Benjamini and Hochberg approach for controlling the false discovery rate (FDR) (Benjamini & Hochberg, 1995). Genes with an adjusted *P*-value of less than 0.05 found by DESeq2 were assigned as differentially expressed. Gene Ontology (GO) enrichment analysis of the DEGs was performed by the DAVID Bioinformatics Resources (https://david.ncifcrf.gov/summary.jsp). We used ToppCluster (https://toppcluster.cchmc.org/) to construct the subcategory network. Heatmaps of gene expression were generated using the Heatmapper (http://www.heatmapper.ca/).

### Quantification of mRNA expression level by real-time PCR

Total RNA was extracted from cells using ISOGEN II (NIPPON GENE, Japan). cDNA was made using 2 µg of total RNA with a SuperScript IV VILO Master Mix (Thermo Fisher Scientific) in accordance with the manufacturer’s instructions. Quantitative PCR (qPCR) was performed using CFX96 Real-Time System (Bio-Rad). Reactions (10 µl each) were prepared in duplicate in Hard-Shell PCR 96-Well Reaction Plate (Bio-Rad). Each reaction contained 0.5 μM of each primer, 10 µl of KAPA SYBR Fast qPCR Master Mix (KAPA Biosystems) and 0.1 ng/µl of cDNA as a template. The expression levels were normalized to UBC (Ubiquitin C) and calculated by the 2^−ΔΔCt^ method. qPCR settings were as follows: initial denaturation at 95 °C for 30 sec, followed by 40 cycles of denaturation at 95 °C for 10 sec, and annealing and extension at 60 °C for 20 sec. The primers used for the qPCR are as described in Table EV1.

### Measurement of cell growth

Cell growth was monitored by a high content screening (HCS) method. Ten thousand cells were seeded per well on 96-well plates (Greiner bio-one #655090) at 14 h before measurements. Prior to imaging, cells were stained with 0.5 μg/mL Hoechst 33258 (Sigma) and 0.15 μM DRAQ7 (BioStatus, #DR71000) to identify live and dead cells, respectively. Images were acquired on a ImageXpress Micro XL High Content Screening (HCS) System (Molecular Devices) equipped with climate control (37 **°**C, 20% CO_2_) using a 10X objective lens. In each condition, cells were assayed in the minimum of 4 fields per well per time point (25 time points with 3 h intervals; total 72 h) were imaged.

### Adipogenic cell differentiation assay

Cells were grown in 6-cm dishes to 80% confluence, and then were cultured in α-MEM (Nacalai tesque, Japan) supplemented with 10% FBS, 1µM dexamethasone (Sigma-Aldrich), 10 µg/ml insulin (Sigma-Aldrich), 100 mM indomethacin (Sigma-Aldrich) and 0.5 mM 3-methyl-1-isobutylxathine (Sigma-Aldrich) for 2 days, followed by incubation for 14 days in maintenance medium (α-MEM, 10 % FBS and 10 µg/ml insulin) (Pittenger *et al*, 1999). Treated cells were fixed and stained with 0.5% oil red O (Sigma-Aldrich) in 60% isopropyl alcohol for 15 min to detect lipid droplets.

### Anchorage-independent assay

Cell transformation was evaluated by measuring anchorage-independent growth (colony formation in soft agar). Base agar is 2 ml of 0.5% Bacto agar (BD Difco) in complete growth medium in each 60-mm dish. Soft agar composed of 2 ml of 0.3% noble agar (BD Difco) in complete growth medium and 1 × 10^3^ cells were .overlaid on base agar. Cells were incubated at 37 °C for 3 weeks, and resulting colonies were counted after staining with 10 % crystal violet in methanol.

### Flow cytometry

To quantify GFP/mCherry fluorescence, cells were analyzed using LSRFortessa flow cytometer (BD Biosciences). BM-MSC were phenotypically characterized by flow cytometry based on the expression of surface markers (CD45 negative, CD73 positive, CD90 positive, and CD105 positive) according to the recommendations of the International Society for Cell & Gene Therapy (ISCT) (Dominici *et al*, 2006). Antibodies used for the staining were all purchased from Biolegend; APC anti-CD45-FITC, APC anti-CD73, APC/Cy7 anti-CD90, and APC anti-CD105. For isotype staining, Cy5-linked goat anti-mouse IgG was used. Cell suspensions were acquired using a LSRFortessa flow cytometer (BD Biosciences), and data were analyzed using FlowJo software (BD Biosciences).

### Immunofluorescence microscopy

Parent and fusion clone cells were seeded onto 15-mm coverslip in 12-well plates, and then were cultured for 24 h. The cells were fixed with 4% paraformaldehyde (Sigma-Aldrich) for 10 min at room temperature and penetrated with Perm/Wash buffer (#554723; BD Biosciences) for 30 min. The cells were blocked with 10% Pierce™ Protein-Free (PBS) Blocking Buffer (#37572; Thermo Fisher Scientific) in PBS for 1 h at room temperature. The cells were immunostained with the specific antibodies overnight at 4°C, washed three times with PBS before incubation with the appropriate secondary antibody and three further washes in PBS. Coverslips were mounted using ProLong™ Gold antifade mountant with DAPI (Thermo Fisher Scientific). Fluorescence was detected using a laser scanning confocal microscope (LSM780; Carl Zeiss). Localization of protein was analyzed using a Zen software (Carl Zeiss).

### Western blotting analysis

The protein extraction and immunoblotting were performed as previous described (Tajima *et al*, 2011). The whole cell extracts were obtained by lysis with RIPA buffer (0.1% sodium dodecyl sulfate (SDS), 1% sodium deoxycholate, 1% Triton-X 100, 150 mM NaCl, and 5 0mM Tris-HCl [pH8.0]) with cOmplete protease inhibitor cocktails (Sigma-Aldrich). We quantified the protein concentrations by microBCA protein assay kit (Thermo Fisher Scientific). RIPA extracts (50 µg of protein) were treated in NuPage lithium dodecyl sulfate (LDS) sample buffer (4X) (Thermo Fisher Scientific) supplemented with 50 mM dithiothreitol (DTT) at 95 °C for 2 min. Proteins were separated by SDS-PAGE on 4-20 % gradient precast gel (EZBiolab Precast Gel, WSHT, Shanghai) at 100 V for 75 min. The separated proteins were transferred on to 0.45 µm PVDF membrane (Merk Millipore, #IPVH00010), followed by blocking with 5 % skim milk in TBS-T. Membranes were incubated with primary antibodies at 4 °C overnight. GAPDH and α-Tubulin served as a loading control (1 : 5000). After wash in 0.1% TBS-T, the membrane were incubated with the appropriate secondary antibodies goat anti-rabbit IgG, peroxidate-linked (1: 5000) or goat anti-mouse IgG, peroxidase-linked (1:5000) at room temperature for 1 h. Peroxidase-linked antibody signal was visualized in Chemi-lumi one Super detection reagents (Nacalai tesque, Japan) using LAS-4000 imaging system (FUJIFILM, Japan) and quantified using Image*J* software.

### Generation of knockout cell lines using CRISPR-Cas9

Gene knockouts were generated by the lentiCRISPRv2 system (Addgene plasmid #83480 and #98291)(Sanjana *et al*, 2014) using the methods described above and specific sgRNA-encoding oligos (Table EV1). Our sgRNA design followed the methods available at Dr. Feng Zhang’s lab (Broad Institute; *https://www.genscript.com/gRNA-database.html?src=pullmenu*). Three different sgRNA sequences were designed for each gene to be knocked out. Briefly, a mixture of equal amounts of guide RNA forward and reverse oligonucleotides were allowed to form duplexes by incubation at 100°C followed by the cooling down process. The duplex was then ligated with lentiCRISPRv2 precut by Bsm BI (New England Biolabs) using T4 DNA ligase (Takara Bio). The ligation mixes were transformed into Stbl3 competent cells and the resultant clones were screened by checking the insert sizes and direct sequencing. Lentiviral particles were produced in 293FT (2 × 10^6^) cells in a 10 cm dish by co-transfection of vectors with lentiCRISPR V2-sgRNA (2 µg), pLP1 (2 µg), pLP2 (2 µg), and pLP/VSVG (2 µg) (clontech) using Polyethylenimine (PEI) Max (Polysciences). Virus containing supernatant was collected 48 h after transfection and cleared through a 0.45-µm PDF filter. Recipient cells were infected overnight with the viral supernatant and then replenished with fresh media. After 48 h, transduced cells were cultured in fresh media containing 1 µg/ml blasticidin S hydrochloride (Fujifilm-Wako pure chemical Corp.) or 250 µg/ml Hygromycin B for 10-12 days. Selected CRISPR/Cas9 Knockout cells were used as bulk populations.

### *In vivo* tumor growth

Procedures for experiments with mice were approved by the Institutional Animal Care Committee of Tokyo Metropolitan Institute of Medical Science, in accordance with the Standards Relating to the Care and Management of Experimental Animals in Japan. 1 × 10^6^ cells resuspended in 0.2 ml of Matrigel (BD Biosciences) were subcutaneously injected into the flank of 8-10 weeks-old female immunodeficient nude mice (BALB/c AJcl-Foxn1^nu^; CLEA Japan Inc.). Tumor dimensions were measured weekly with serial calipers and volumes were estimated by the modified ellipsoidal formula (Tomayko & Reynolds, 1989); tumor volume = length x width^2^/2. Mice were sacrificed at 12 week or when tumor volume approached 3,000mm^3^.

---

**Figure EV1.**
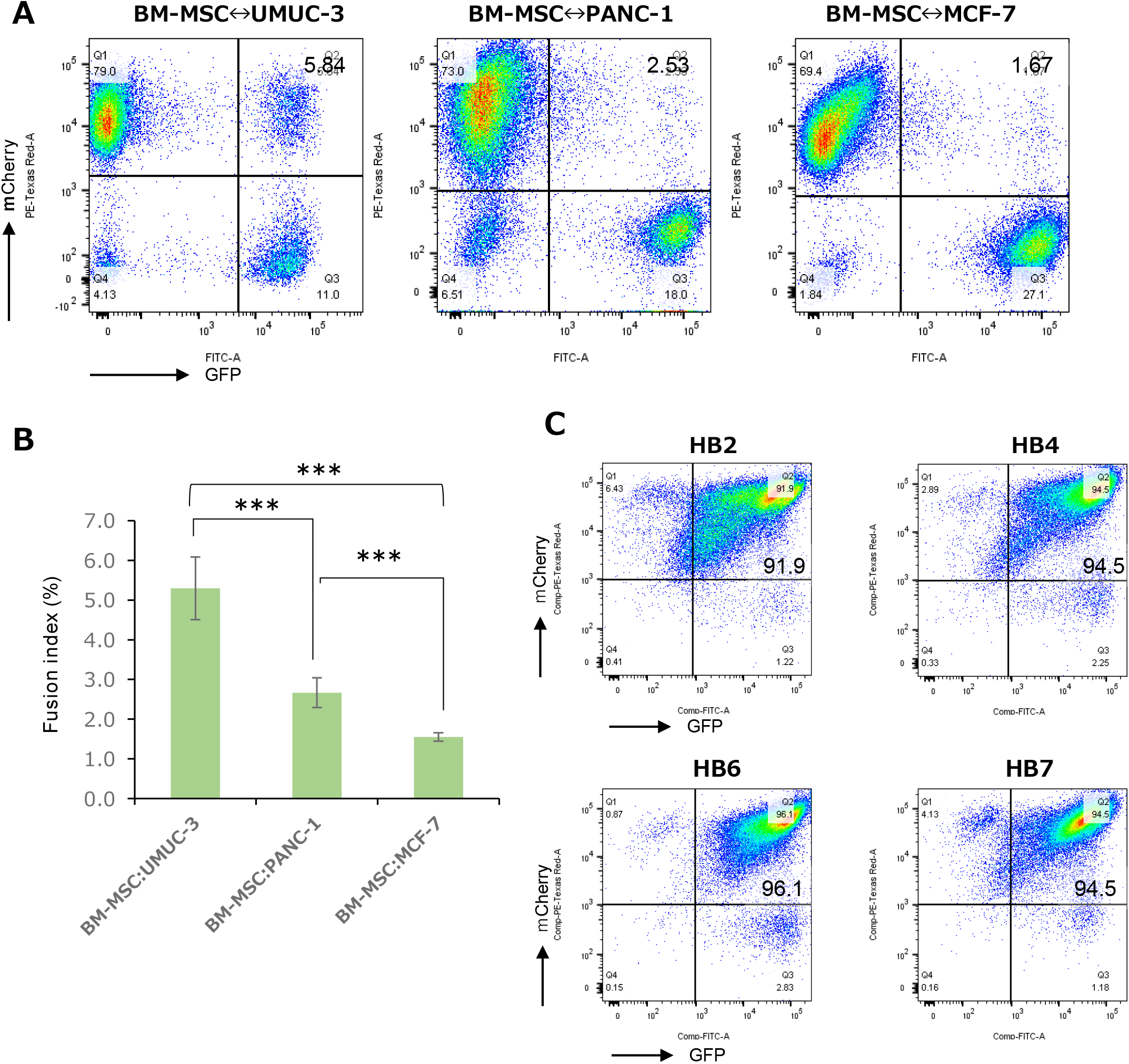
Cancer cells fuse spontaneously with BM-MSC. **A**. Flow cytometry analysis of GFP (FITC-A channel) and mCherry (PE-Texas Red-A channel) after coculture of GFP-positive BM-MSC and each of mCherry-positive three tumor cell lines UMUC-3, PANC-1, and MCF-7 for 2 days. The fractions of the GFP and mCherry double positive fusion cells are shown in upper right quadrants of each graph. **B**. Fusion rates calculated by the data of flow cytometry. Each value represents the mean (± SD) from at least three independent experiments. *** indicates *P* < 0.001, using two-tailed Student’s *t*-test. **C**. Flow cytometry analysis of GFP (FITC-A channel) and mCherry (PE-Texas Red-A channel) in the fusion cells (HB2, HB4, HB6, and HB7). More than 90 % of the cells are GFP and mCherry double positive.

**Figure EV2.**
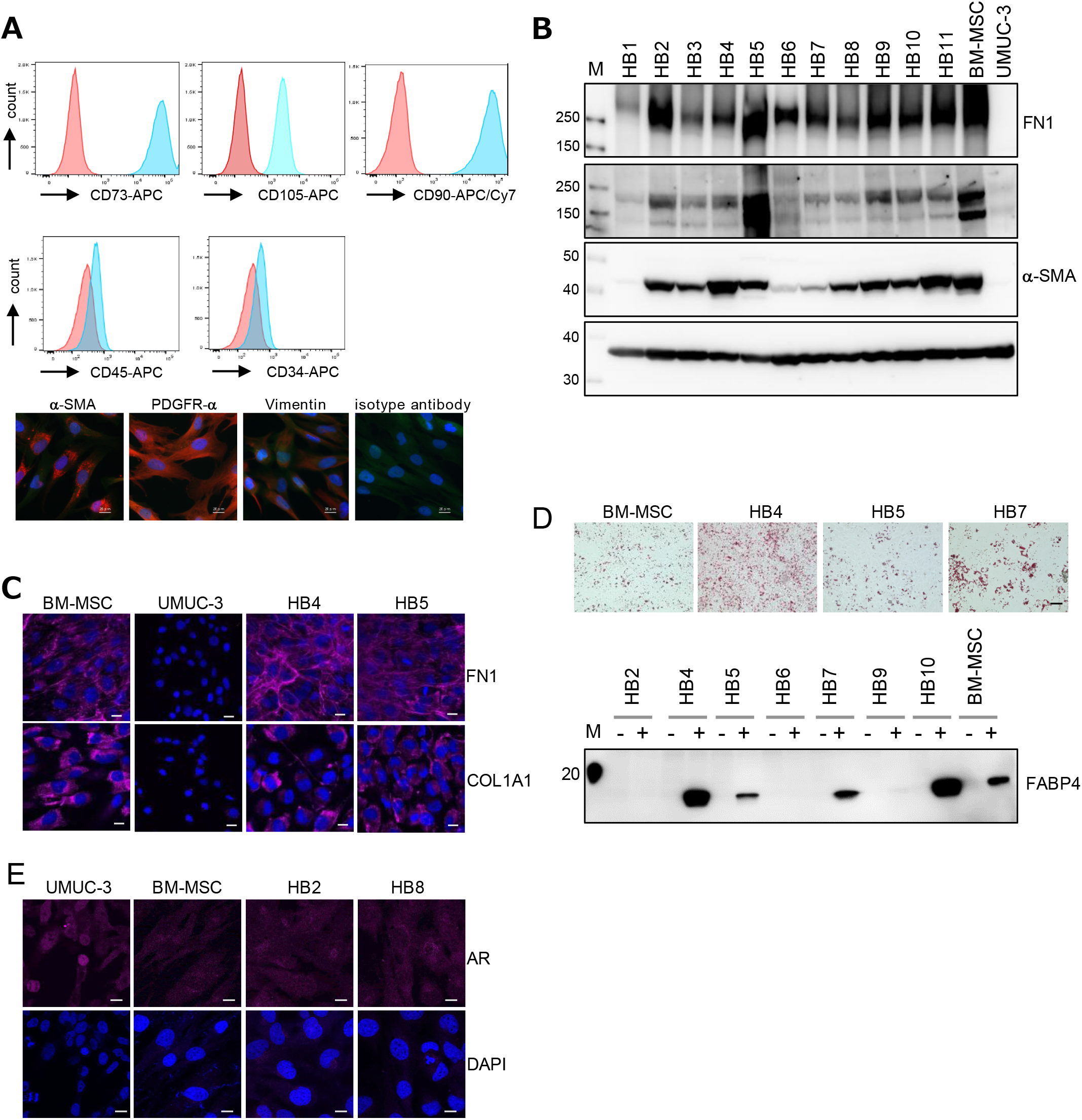
Fusion cells display mesenchymal stem cells properties. **A**. Upper, Cell-surface marker profiles of BM-MSC cells determined by flow cytometry using antibodies against CD34, CD45, CD73, CD90, and CD105; blue and red represent specific antibodies and isotype controls, respectively. Lower, BM-MSC cells were immunostained with antibodies against α-smooth muscle actin (α-SMA), platelet-derived growth factor receptor-α (PDGFR-α), and vimentin, markers of MSC-derived myofibroblasts. Bar indicate 20 µm. **B**. Western blot analysis of proteins indicated in parental (BM-MSC and UMUC-3) and 11 fusion (from HB1 to HB11) cells. **C**. Immunostaining of FN1 and COL1A1 in parental and fusion cells (HB4 and HB5). Scale bar indicate 20 µm. **D**. Upper, Adipogenic differentiation was assayed by Oil Red O staining for lipid vesicle formation in BM-MSC, HB4, HB5 and HB7 grown in the induction medium. Scale bar indicate 100 µm. Lower, Western blot analysis of fatty acid binding protein 4 (FABP4), a marker of adipocytes, in BM-MSC, HB4, HB5, HB6, HB7, HB9 and HB10 grown in induction medium (+) or without induction (-). **E**. Immunostaining of androgen receptor (AR), a marker of urogenic tumors, in parental and fusion cells (HB4 and HB5). Scale bar indicate 50 µm.

## Acknowledgments

We thank Yoshinobu Iguchi and Kazunari Sekiyama for preparation of cryosections. We also thank Dr. Daisuke Yamane for the insightful discussion and comments. We thank Hiroyuki Sasanuma for critical reading of the manuscript. This work was supported by JSPS KAKENHI Grant-in-Aid for Scientific Research (A) Grant Number 20H00463 (to HM) and by KAKENHI Grant-in-Aid for Scientific Research (C) Grant Number 18K07220 (to YT) and KAKENHI Grant-in-Aid for Challenging Exploratory Research Grant Number 16K15262 (to YT), respectively.

## Author contributions

Conceptulization: YT and HM; Methodology and Investigation: YT; Writing and Visualization: YT and HM; Supervision: HM and FS; Funding Acquisition and Project Administration: HM.

---

**Table EV1:**
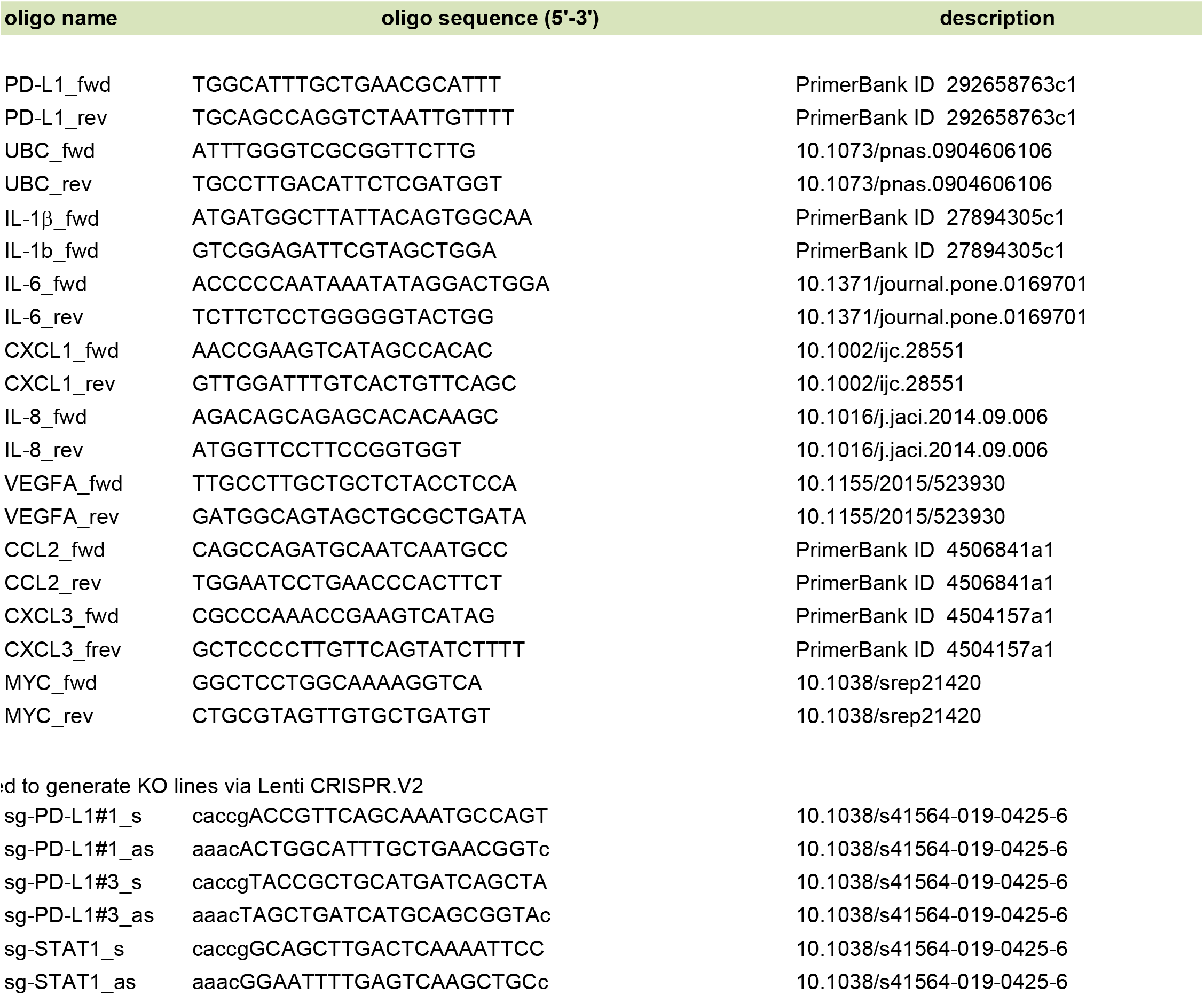

